# Languages, evolution and statistics: human sound systems were shaped by changes in bite configuration. *Response to Tarasov & Uyeda (2020)*

**DOI:** 10.1101/2020.02.27.965400

**Authors:** Damián E. Blasi, Steven Moran, Scott R. Moisik, Paul Widmer, Dan Dediu, Balthasar Bickel

## Abstract

In Blasi et al. (2019) we have shown, through a series of statistical analyses and models, that human sound systems have been affected by a transition in bite configuration starting from the Neolithic. Tarasov and Uyeda (2020) (henceforth T&U) raise a number of observations in relation to our article. We appreciate T&U’s engagement with our work and their sharing of the code and data of the analyses reported. In brief, their technical comment involves five analyses:

1. Binomial Causal Models (BCM)
2. Linear Regression of across-area variation in labiodentals and subsistence
3. Predictive Posterior Simulations (PPS)
4. Poisson Linear Regression (PLR): model comparison
5. Phylogenetic Analyses

In what follows, we show that the discrepancies they report between our findings and theirs are due mostly to ill-specified models, weak (or missing) statistical evidence, and a misinterpretation of our results. After these issues are addressed, we conclude that T&U’s claims do not hold.

## 1. Binomial Causal Models (BCM)

The first study focuses on the association between labiodentals and subsistence within areas. Their argument starts with the point that area might be causally responsible for the development of certain types of subsistence and, at the same time, through the historical relations that hold between languages, for the presence of labiodentals within them. Then they conjecture that the very association between labiodentals and subsistence might be accidental, as it would occur when no adequate controls are taken into account. Figure R1 is a graphic display of the causal dependencies, where solid arrows imply causal relations between the variables expressed in the nodes.

**Figure R1:**
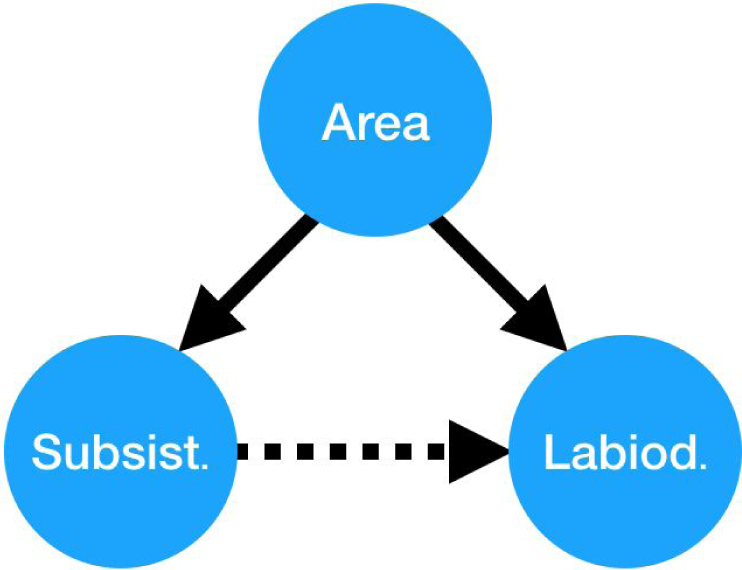
Graphic display of causal dependencies. Solid arrows represent causal connections, and the dotted one, a spurious one.

In their study, T&U implement simple mono-parametric Bayesian binomial regression models aimed at accounting simultaneously for (1) the number of hunter-gatherer societies, (2) the number of hunter-gatherer societies that have labiodentals, and (3) the number of food producing societies that have labiodentals. They statistically contrast two models: (A) a model where the parameters for (2) and (3) are different (the independent model), and (B) a model where they are identical (the dependent model). However, we note several substantial issues with their models and their interpretation of our study:

- Minor issues in interpretation, data coding, and data selection
- Marginal vs. conditional models
- No causal guarantees
- Convergence issues in the MCMC chains
- Insufficient Bayesian power
- No genealogical controls

We discuss these in turn.

### 1.1. Minor issues in interpretation, data coding, and data selection

There are three minor observations about the test and its evaluation that, while not fatal to the inference, still merit close observation and raise concerns.

First, we locate the causal mechanism in the *biomechanics* of speech production, its effects on accidental errors, and the probability that these errors stabilize in a population of speakers, ultimately leading to language change. We use the subsistence and historical data as *proxies* for differences in bite between speaker groups, and *not as direct causal factors* for the emergence of labiodentals.

Second, it is worth mentioning that instead of evaluating the response variable we associate with our claim (the *number* of different labiodentals in the language) T&U resort --- presumably due to statistical convenience --- to the analysis of *presence/absence* of labiodentals. While the difference between the two might not be large in practice, the rationale behind our choice resides in the fact that one marginal labiodental might exist in a given language as the result of a handful of borrowed words, whereas two --- or more than two --- contrastive labiodentals in a language more strongly support the notion that labiodentals are an *organic part* of the system, independently from recent and sporadic contact.

Third, it is interesting to note that T&U isolate the Atlantic-Congo family from Africa and repeat the test with just this family, presumably finding no association after the data are filtered. The only expressed motivation is that “the majority of languages in Africa belong to the Atlantic-Congo family”, which strikes us as a symptom of a biased evaluation: for instance, they do not repeat the analysis after removing the *Austronesian* family from the Pacific (where no association is found), even if it is true that the majority of languages in the Pacific belong to that one family (so, a similar argument as for Africa should apply). Subsetting the data conditioning on the result of a test can easily lead to erroneous and biased inferences, and opens the door to evidence fishing

### 1.2. Marginal vs. conditional models

Before discussing the details of the T&U model and its evaluation, it is important to spell out their interpretation of what the observed pattern of association between labiodentals and subsistence should be according to our (and Hockett’s) hypothesis.

*“If Hockett’s hypothesis is true, then the dependent model, indicating causality between subsistence and labiodentals*, ***is expected to be selected across all regions****” (T&U, P2, emphasis ours)*

In other words, they expect that each of the areas in our study should provide evidence of the association between labiodentals and subsistence *after* aggregating all other factors shaping the presence of labiodentals (i.e. language history, overall inventory size, etc.).

This is a *marginal* reading of the hypothesis, whereas our claim (and Hockett’s) is *conditional*: *once all other effects have been taken into account*, it should be possible to detect that food producing societies have more labiodentals than hunter-gatherers. Formally:

T&U claim: labiodentals ∼ subsistence

Our claim: labiodentals ∼ subsistence | covariates

The marginal reading is much stronger than the conditional one, and in general we expect it to be false for almost all claims about the adaptation of human behavior to external factors across the world, given the complexities of human behaviour and culture, and the many factors --- some only due to historical contingencies --- that affect them. Multiple transmission biases affecting culture, coupled with competing adaptive pressures, might bias human societies to adopt a trait that would be generally dispreferred, but not when taking into account all these conditions. The covariates we consider in the conditional reading of the hypothesis are well-known to mask universal preferences in language (Dryer 1989; Jaeger et al. 2011; Bickel 2011).

Furthermore, our own modelling shows that the *marginal* hypothesis does not hold: upon inspecting the random intercepts assigned to different areas in the full models, we find that the extent of variation in areal biases towards vs against labiodentals is comparable to the main effect of subsistence on labiodentals (and, importantly, the majority of these estimates are comparable across both datasets analyzed). The details are shown in Figure R2.

**Figure R2:**
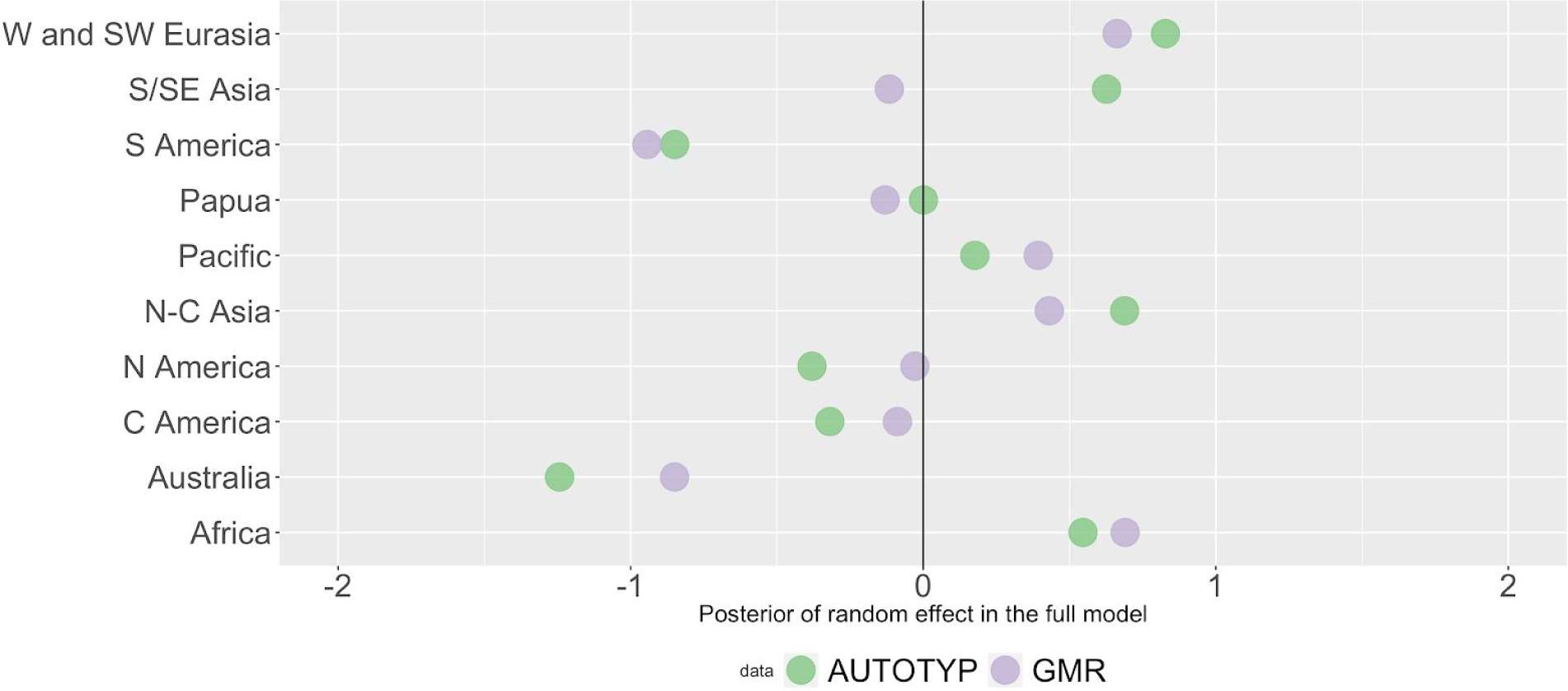
Mean of the posterior of random effects in the full model by area, based on our original results

A smaller, yet still detectable effect, is represented by the effect of language family random intercepts, as shown in Figure R3.

**Figure R3:**
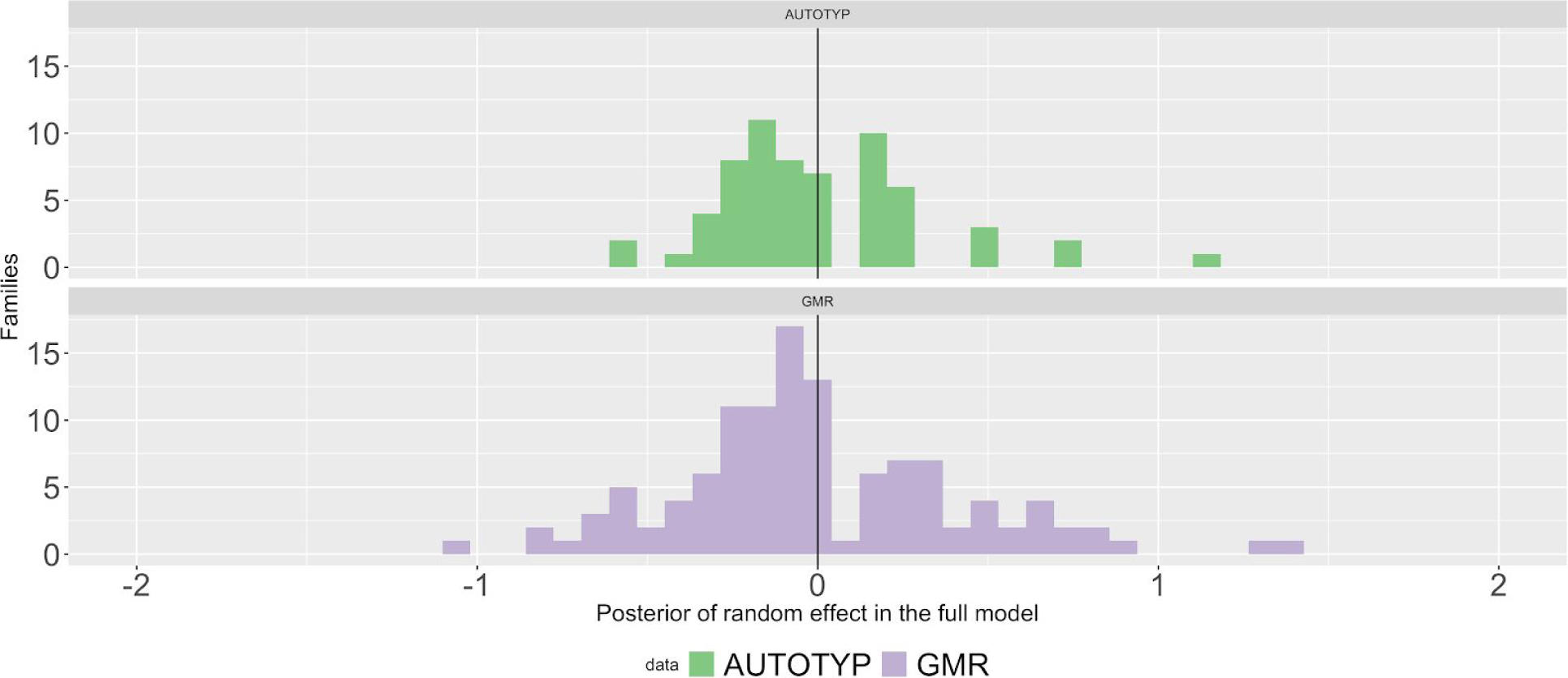
Mean of the posterior of the random effects in the full model by language family: again, we see that there is variation across families, but is relatively smaller than for areas (Figure R2).

In sum, our results support the notion that individual families and areas might not follow the overall trend. Crucially, we spell this out clearly in the main text (Blasi et al. 2019):

*The mean posterior coefficient of subsistence on the presence of labiodentals, while displaying a wide posterior, is comparable in magnitude to the characteristic differences between linguistic families and areas—in other words, the models suggest that differences in subsistence have as substantial an impact on labiodentals as do the differences among families or geographical areas.*

Thus, even before discussing the statistical issues with T&U’s analyses, we must note that they misinterpreted our hypothesis, making, thus, their claims irrelevant. Nevertheless, it is important to note that T&U’s own model for the whole dataset (denoted as “Whole World” in their Table 1) *does* yield support for the strong, marginal hypothesis.

### 1.3. No causal guarantees

T&U begin their technical comment by noting that:

*“While regression models are powerful statistical tools for inferring correlation between variables, it is of course true that correlation does not mean causation (…) Regression analysis alone cannot disentangle these different types of dependencies, and can often result in overstating the evidence for a causal relationship.” (P1-2)*

We wholeheartedly agree, and that is *precisely why* the correlation analysis we produce is but one line of argumentation among a larger set of mutually reinforcing studies involving: (a) the simulation of speech articulation, (b) philological studies on specific world areas, (c) a review of the state-of-the-art of the paleoanthropological evidence, and (d) an in-depth phylogenetic analysis on a well-studied language family (Indo-European).

There are developments in the field of causal inference with observational data where a stricter set of conditions on statistical models yield causal guarantees of different types. Perhaps the two best-known approaches for causal inference with observational data are the *do-calculus* and the language of *DAGs* (Pearl (2009), applied to linguistic inference in Blasi (2018) and models of *additive white noise*. Thus, after reading the previously highlighted line in T&U’s response, we expected an approach of this kind. Surprisingly, T&U’s proposal to fix the alleged causal deficit of our models is an even simpler regression model.

Barring the difference between ours and T&U’s response variable (pointed out above), we could express our models as a triad:

1. labiodentals ∼ intercept + subsistence + interaction(subsistence,area) + interaction(subsistence,family) + covariates
2. labiodentals ∼ intercept + interaction(subsistence,area) + interaction(subsistence,family) + covariates
3. labiodentals ∼ intercept + covariates,

where the left hand side stands for the response variable (labiodentals) and the right hand side includes the main effects and interactions.

For each of our datasets (*AUTOTYP* and *GMR*) and each of our three conditions (*all areas, without Australia, fricatives instead of all segments*) we produce three Bayesian models that allow us to perform state-of-the-art model evaluation leading to a decision on the relative performance of subsistence as a main effect and in interaction with selected covariates.

T&U’s model, using the same notation, tests *within individual areas*:

1. labiodentals ∼ intercept + subsistence
2. labiodentals ∼ intercept

Note that this model is a marginal model (discussed above) and that it violates iid assumptions (discussed below). We fail to see in which conceivable way their much simpler regression analysis and model selection, has any causal guarantee that our models do not.

Furthermore, they test:

(3) number of hunter-gatherer societies ∼ intercept

but the role of this model is unclear, and it does not play any role in the evaluation of the role of subsistence on labiodentals. Given the uninformative prior T&U use, the intercept of model (3) will trivially be the ratio of all hunter-gatherer societies over the total number of societies in such areas. Unless one wanted to obtain a measure of uncertainty of that estimate, there is no need of deploying this model for estimating the binomial parameter. But the posterior uncertainty never shows up in their analyses.

### 1.4. Convergence issues in the MCMC chains

After reproducing T&U’s analysis using their provided code, we encountered that several models yielded issues of convergence of the kind of post-warm-up divergent transitions in Stan (a probabilistic programming language for statistical inference). Stan’s manual highlights the importance of such warnings:

*Stan uses Hamiltonian Monte Carlo (HMC) to explore the target distribution — the posterior defined by a Stan program + data — by simulating the evolution of a *Hamiltonian* system. In order to approximate the exact solution of the Hamiltonian dynamics we need to choose a step size governing how far we move each time we evolve the system forward. That is, the step size controls the resolution of the sampler.*

*Unfortunately, for particularly hard problems there are features of the target distribution that are too small for this resolution. Consequently the sampler misses those features and returns biased estimates. Fortunately, this mismatch of scales manifests as divergences which provide a practical diagnostic.*

And later,

***Since the validity of the estimates is not guaranteed if there are post-warmup divergences***, *the slower sampling is a minor cost.*

We think that the issues emerge from problems that have to do with the absence of data for estimating the models (to which we return to below); the fact that such numerical issues are not discussed in T&U is problematic.

### 1.5. Insufficient Bayesian power

The marginal hypothesis put forth by T&U requires the association to be present in every area, but it does not entail that there will be enough statistical power so as to be able to leverage the association.

After T&U note support for the marginal model in Africa and NC Asia, they observe that “in the remaining eight areas, the independent model exhibits better or similar fit, thus contradicting Hockett’s hypothesis”.

However, T&U do not conduct a Bayesian power analysis nor do they spell out what would be the effect size that would be consistent with the marginal hypothesis. Yet even under this underspecification, it is straightforward to see that many individual areas have very little data, as displayed in Table R1.

Figure R4 display this heavily skewed distribution on a map.

**Figure R4:**
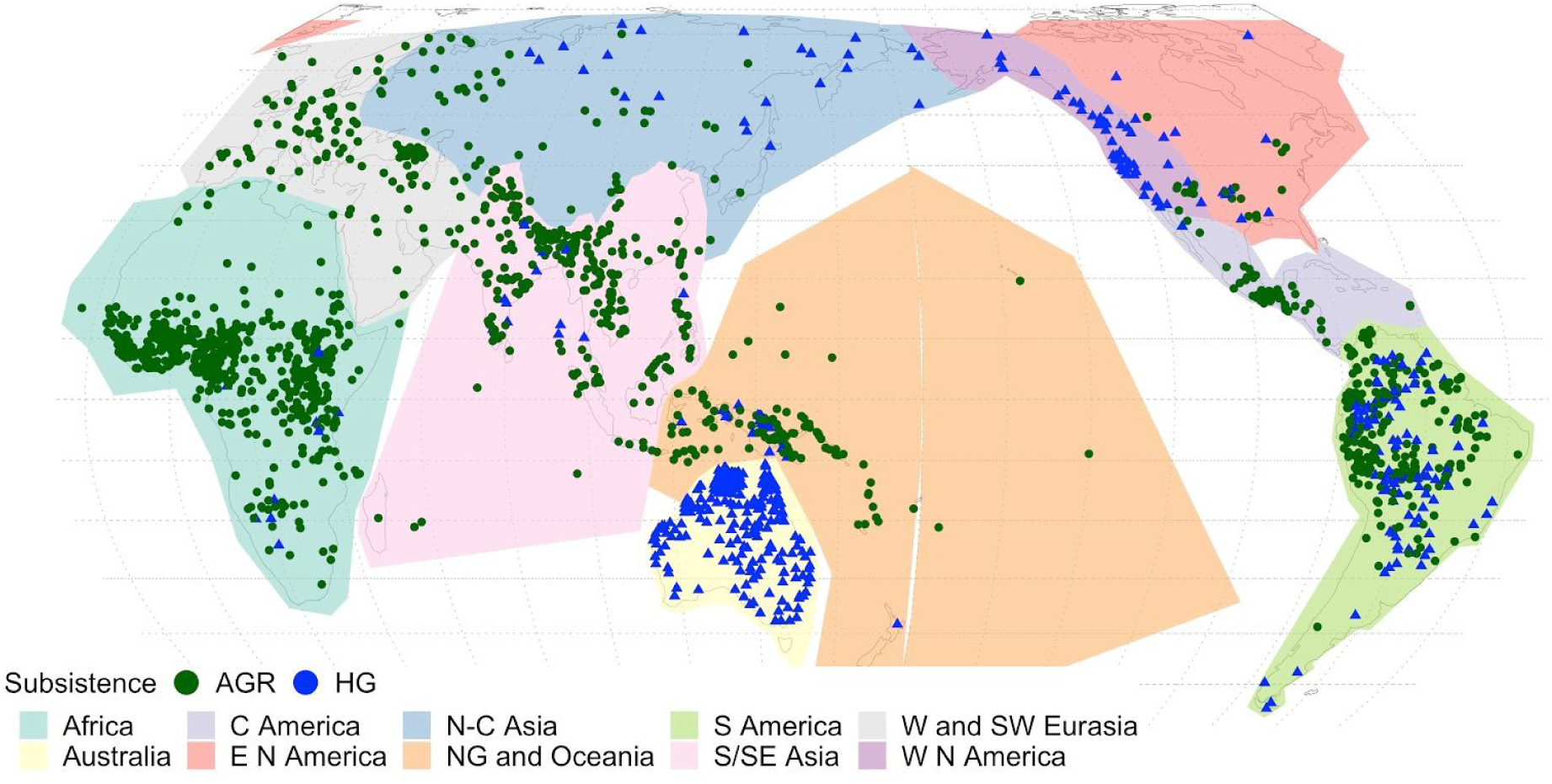
Worldwide distribution of subsistence by area.

**Table R1:** Bayesian power analysis of GMR and AUTOTYP data. Red rows highlight cases that are extremely problematic due to low degree of variation.

| Area | Data | HG | FP |
| --- | --- | --- | --- |
| Africa | GMR | 5 | 696 |
|  | AUTOTYP | 8 | 15 |
| Papua | GMR | 26 | 64 |
|  | AUTOTYP | 5 | 20 |
| S America | GMR | 100 | 226 |
|  | AUTOTYP | 42 | 48 |
| W and SW Eurasia | GMR | 0 | 110 |
|  | AUTOTYP | 0 | 18 |
| C America | GMR | 1 | 52 |
|  | AUTOTYP | 1 | 11 |
| S/SE Asia | GMR | 13 | 279 |
|  | AUTOTYP | 7 | 19 |
| N America | GMR | 67 | 27 |
|  | AUTOTYP | 64 | 8 |
| Pacific | GMR | 2 | 53 |
|  | AUTOTYP | 2 | 1 |
| Australia | GMR | 242 | 0 |
|  | AUTOTYP | 107 | 0 |
| N-C Asia | GMR | 20 | 46 |
|  | AUTOTYP | 21 | 10 |

Most areas are strongly biased towards one or another type of subsistence. Worryingly, in four of these areas there is a whole subsistence category with two or less observations. Given the nature of subsistence as a proxy for bite configuration, the coding inconsistencies in subsistence type that emerge from any ethnographic record and the extremely low power implied by such small samples, it is clear that such models are uninformative, at best, and they are, in fact, very likely to be misleading.

To assess this formally, we performed a Bayesian power analysis by taking the marginal effects on the data. We use as representative values of the binomial parameters for each group (θ_HG_ and θ_FP_) the raw conditional probabilities as taken in the whole datasets. Thus θ_X_ is the proportion of all languages in group X that have labiodentals. HG display across the board around θ_HG_=0.25 and FP θ_FP_=0.55. This is a substantial difference between the two groups. We then used T&U’s code to produce artificial data simulated out of the total number of HG and FP languages in each area while assuming the observed parameters of θ_HG_=.25 and θ_FP_=.55 respectively. We run the simulations N=20 times for each area, and then collect the fraction of times |ln(BF)|>1.1, which, according to standard guidelines, corresponds to the minimum threshold for determining any evidence against or in favour of a given hypothesis. The results are given in Table R2.

**Table R2:** Results of Bayesian power analysis by area. Cells display the proportion of simulations in which there is support for the hypothesis.

| Area | Bayesian power |  |
| --- | --- | --- |
|  | AUTOTYP | GMR |
| Africa | 0.05 | 0.2 |
| Papua | 0 | 0.65 |
| S America | 0.6 | 0.85 |
| W and SW Eurasia | 0 | 0 |
| C America | 0.1 | 0 |
| S/SE Asia | 0 | 0.4 |
| N America | 0 | 0.55 |
| Pacific | 0 | 0 |
| Australia | 0 | 0 |
| N-C Asia | 0.05 | 0.4 |

The results are very clear: even for testing the (wrong) marginal model, there is simply no data. If one follows the usual principle of guaranteeing a minimum of 0.8 power for a statistical evaluation, they will only be left with the test for South America exclusively in the GMR data (which is the area with the most diversity in the data; see Figure R4)

In other words, T&U conflate absence of evidence with evidence of absence.

### 1.6. No phylogenetic controls

Lastly, T&U’s model does not include any controls for phylogenetic dependencies between languages. Control for phylogenetic autocorrelation has been a standard concern in language science since the late 1980s (Dryer 1989; Jaeger et al. 2011). Hence their model entails that, conditioning on area and subsistence, the presence of labiodentals within any given area is independent from their presence in other languages (or in other words, it violates the assumption of observations being independent and identically distributed (iid). Given that we established in our study that vertical transmission (language genealogy) does indeed play an important role in the presence of labiodentals (see Figure R3 above), this omission is extremely problematic.

## 2. Linear Regression of across-area variation in labiodentals and subsistence

This model still suffers from some of the issues outlined for the previous study (e.g. lack of controls for shared history), but since the results support the marginal model which builds on a different hypothesis than ours our Hockett’s, we see no reason to discuss it.

## 3. Predictive Posterior Simulations (PPS)

T&U compare the performance of a number of Poisson regression models using five possible different variables as predictors (areas, distance Out-of-Africa, number of non-labiodental fricatives, number of non-labiodental phonemes, subsistence) and a model with a single common intercept.

Our criticism can be divided into two: technical objections and theoretical ones, the later focused on the assumed model that T&U utilize as a way of introducing Out-of-Africa distance as a viable variable in the model.

### 3.1. Technical objections

Once more, T&U cast a marginal model whereas our hypothesis entails a conditional model. The criticism is identical to the one we discussed in relation to T&U’s Study 1 (see above). We do not embrace a marginal model of association between subsistence and labiodentals presence, hence evaluating the performance of models built on individual predictors is orthogonal to our claim. Area, language family and/or number of non-labiodental segments could display a larger explanatory power individually and yet it would be possible to identify a conditional effect of subsistence (as it is indeed the case in our case).

### 3.2. Theoretical objections

In their rebuttal, T&U assume that distance-from-Africa is a widely accepted confound. They assume at face value that “phonemic diversity” (defined as the number of phonemes in a language) decreases with a language’s distance from Africa due to a serial founder effect as reported by Atkinson (2011). Atkinson’s proposal is certainly intriguing, as it claims to have found a linguistic signal that traces language’s origins back to Africa, some 50-70kya. This time-depth is much further than what the historical comparative method can reconstruct in terms of proto-languages, which is limited to a time depth of roughly 10kya (e.g., Renfrew et al. (2000)). As such, Atkinson’s claim goes against current linguistic theory about rates of language change.

Also, the time-depth of Atkinson’s model is very different from our focus on changes since the Neolithic. Thus, an increase in labiodental sounds due to changes in human bite configuration from access to softer diets would have occurred much later after the expansion of humans across the globe and involved many technological and cultural changes. Given this, Atkinson’s and our (or Hockett’s) hypotheses cannot be directly compared, without any further control of time depths.

These discrepancies notwithstanding, it is important to note that Atkinson’s analysis has not held up to scientific scrutiny. See the technical comments and reanalyses in *Science* (Cysouw, *Dediu, and Moran 2012; Van Tuyl and Pereltsvaig 2012; Wang et al. 2012) and in Linguistic Typology* (Bybee 2011; Dahl 2011; Donohue and Nichols 2011; Jaeger et al. 2011; Maddieson et al. 2011; Pericliev 2011; Ringe 2011; Ross and Donohue 2011; Sproat 2011). In summary, the main criticisms and issues are:

1. Atkinson’s problematic definition of phonemic diversity
2. There is no positive correlation between population size and phoneme inventory size
3. Contradictory points of language’s origin
4. The processes involved in how phoneme inventories change
5. No theoretical or empirical support for a serial founder model for language

We explain these in turn.

#### 3.2.1. Atkinson’s problematic definition of phonemic diversity

Several studies show that Atkinson’s definition of phonemic diversity is not a valid measure of phonological complexity (Sproat 2011; Maddieson et al. 2011; Cysouw, Dediu, and Moran 2012; Wang et al. 2012). Atkinson’s linguistic data source, the *World Atlas of Language Structures* (Haspelmath et al. 2008) includes summaries of consonant, vowel, and tone inventory counts from roughly 10% of the world’s languages (Maddieson 2011). In these databases, phoneme inventory sizes are simplified by binning total phoneme counts into three-to-five levels (small, medium, large) for consonant, vowel, and tone inventories.

Wang et al. (2012) argue that this simplification of binning phoneme inventory counts into levels is questionable and they find that Atkinson’s results are only supported when phonemic diversity is binned. They note that:

- Consonant inventories vary in size from 10 to more than 80, but discretizing this range into five levels obscures the phonological complexity of consonant inventories.
- For vowel inventories, the lower limit of seven vowels eliminates the difference between diversity in African and Eurasian languages.
- Binning phoneme inventory counts ignores all phonetic features of vowel inventories, including distinctions, such as length, nasalization, and diphthongization, which do not introduce single phonemes in a language but rather cause vowel inventory counts to double, triple or even quadruple in size (see discussion below).
- The reduction of tones into bins is particularly problematic for Atkinson’s analysis because the simplification of tone obscures the fact that most languages in Africa have less than four tones (on average three), whereas languages in Asia have generally more than four. Therefore, in WALS tone diversity between the two continents is obfuscated, and Wang et al. (2012) find that when using the raw data of phoneme counts for 510 languages that the highest diversity for tone is in Asia and not Africa, suggesting that WALS binned data bias the results.

Cysouw et al. (2012) also show that binning phoneme inventory counts without any sensitivity analysis of the thresholds goes against current best practices in statistics and may bias the results. And from a theoretical perspective, Ross & Donohue (2011) note that Atkinson conflates phonological features into a single dimension, which collapses the various independent dimensions of phonological complexity.

Maddieson et al. (2011) state that the label “phonemic diversity”, as used by Atkinson (2011), is misleading because:

1. It does not consider each language’s full set of phonemic and prosodic contrasts
2. It does not take into account the internal diversity of each phoneme set.

They provide an example of how the basic vowel inventory as described in WALS (also noted by Wang et al. (2012)) hides the contrast among matching vowels that differ in terms of length, nasalization, voice quality, etc. For example, Maddieson et al. (2011) note that Navajo (North America) has four basic vowel qualities /i, e, a, o/, which can be short or long, and oral or nasal. This results in 16 vowel phonemes, which is coded in WALS as a “small” vowel inventory. This is in contrast to, for example, Igbo (West Africa), which has eight basic vowel qualities, and thus is coded by WALS as having a “large” vowel inventor, even though it has half the number of vowel contrasts as Navajo. Given the coding of vowel inventories in WALS, the Navajo vs. Igbo contrast is far from an exception, but it is emblematic of languages with vowel quantity distinctions across the board.

Liberman (2011) points out that it is problematic that consonant, vowel, and tone inventory counts are binned and then weighted equally in terms of diversity, because their contribution to syllable and word diversity are radically different. As such, Atkinson’s definition of phonemic diversity gives excessive weight to certain features: for example, he suggests that losing a single tone may reduce the total phonemic diversity by as much as ten consonants. And although the loss of a tonal distinction would create syllable pairs that are no longer linguistically contrastive, the loss of a consonant means unequivalent reduction of phonemic variants due to its different potential places within a syllable. In other words, losing a contrastive tone results in the collapse of contrastive pitch on syllables or words, but their basic structure remains the same. On the other hand, when a language loses a contrastive consonant, it affects the language’s lexicon in more ways because consonants my appear in word initial, medial, or final positions.

Van Tuyl & Pereltsvaig (2012) further argue that assigning numerical values to WALS categories is also questionable because the ratios are arbitrary (e.g. each vowel is worth 2.6 consonants, but each tone is 5.7 consonants). This means that in Atkinson’s analysis, consonant inventories play a relatively smaller role in the regression versus distance. Van Tuyl & Pereltsvaig (2012) note that tone dominates the correlation, with tones (59%), vowels (40%), and consonants (1%) making up the African regression slope.

Maddieson et al. (2011) raise a broader issue in that Atkinson’s notion of phonemic diversity is neither a measure of the size of a phoneme inventory nor of the phonotactic possibilities of the languages in the sample. A proper diversity measure should concern the number of distinctions, but phoneme inventory diversity has more to do with an inventory’s contents than its size. Hence, as noted by Sproat (2011), the notion of phonemic diversity requires a significant amount of thought. For example, a language with seven consonants at seven places of articulation is arguably more diverse than a language with seven consonants at three places of articulation. There is not only diversity in phonological features (features themselves are an area of intense linguistic debate), but also diversity in the possible phoneme combinations within a language, i.e. its phonotactics. Hence, Donohue & Nichols (2011), Maddieson et al. (2011), and Wichmann et al. (2011) state that measures of phonological complexity, such as phonotactics and word length, provide a truer measure of diversity. Wichmann et al. (2011) evaluate Atkinson’s results with a different database and find that mean word length is significant, while Maddieson et al. (2011) also assess Atkinson’s claims and note that the number of phonemes is not a true measure of phonological complexity because consonant, vowel, and tone inventories have different effects.

The coarse simplification of languages’ detailed phonological repertoires also obscures the theoretical debates on how to analyze specific sounds in specific languages. For example, should a dually-articulated consonant or diphthong be considered a single phoneme or a sequence of two sounds? Bybee (2011) and Van Tuyl & Pereltsvaig (2012) point out that phonemes are a unit of analysis and different analyses yield very different phoneme inventory sizes (examples are given in Donohue & Nichols (2011), Maddieson et al. (2011), and Rice (2011)). Hence, there are significant variations and uncertainties in the phoneme inventory sizes reported in different sources.^1^

Furthermore, Atkinson’s data are summaries of UPSID (Maddieson 1984) as reported in WALS, but, crucially, Cysouw et al. (2012) show that phoneme inventory size in WALS is imperfectly correlated with the number of phonemes reported in UPSID. This leads to a strong bias in the geographic patterning in Western Africa in Atkinson’s results.

The size of a phoneme inventory is also the outcome of the sound changes that have occurred in that particular language, which results in the gain or loss of phonemes. The relation between sound changes and the number of phonemes in an inventory is indirect (Bybee 2011). Sound change typically affects natural classes of phonemes, and not single phonemes (except in the relatively rare cases of borrowing (Ringe 2011).

#### 3.2.2. There is no positive correlation between population size and phoneme inventory size

In Atkinson’s (2011) view, small populations left Africa and founded new colonies, in which phonemic diversity decreased due to a sampling effect akin to what happens in population genetics. These daughter populations never regained their phonemic diversity, but instead established, in turn, new colonies. Each new colony then has a reduced phonemic diversity, and so on, which ultimately lead to the smallest (on average) phoneme inventories of the languages in South America and Oceania, which are at the furthest points of human migration from Africa.

Atkinson’s model depends crucially on the observation reported by Hay & Bauer (2007), also replicated by Atkinson (2011), that there is a positive correlation between the size of a phoneme inventory and the number of people who speak that language. In other words, larger speaker populations purportedly have larger phoneme inventories.

Furthermore, in our original paper we included the total count of segments among the covariates and showed that the effect of subsistence on labiodental counts persist *even after controlling for the number of non-labiodental segments*. This already casts doubts on T&U’s claim.

These observations notwithstanding, it is worth assessing the hypothesis that phoneme inventory size correlates with population size. The correlation goes back to an observation made by Trudgill (2004), who describes the loss of phonemes during the eastward expansion of Austronesian across the Pacific.^2^ Trudgill did not turn his observation into a general principle regarding phoneme loss and migrations. In fact, he states that he does not believe that migration leads to a reduction in phoneme inventory size (Trudgill 2011). Instead, Trudgill suggests that various social factors, in combination, can influence a speaker community’s phoneme inventory size, such that:

- Large communities sizes favor medium size phoneme inventories
- Small community sizes favor either small or large phoneme inventories

Trudgill maintains that one should not expect to find a meaningful correlation between phoneme inventory size and population size given what is known about phonological change.

Pericliev (2004) tests the observations by Trudgill (2004) and whether there is a positive correlation between speaker community size and phoneme inventory size using the UPSID sample of 451 languages (Maddieson and Precoda 1990) together with modern day population figures from the Ethnologue. Pericliev finds no correlation between the size of a community speaking a language and the size of the phonological inventory of that language.

Hay & Bauer (2007), however, do report a statistically significant positive correlation between population size and phoneme inventory size when using a relatively small sample of 216 languages (Bauer 2007).^3^ The authors are reluctant to offer an explanation for this positive correlation and instead they urge caution in interpreting their findings.

A contrasting finding is reported by Donohue & Nichols (2011), who use a much larger sample of languages with greater worldwide genealogical and areal coverage. They attempt to replicate Atkinson’s findings regarding population size, but they find no correlation.

However, both Hay & Bauer (2007) and Donohue & Nichols (2011) are flawed in terms of statistical approach. Both studies use statistical models that are inappropriate for capturing the non-independence of language families. Moran et al. (2012) remedy this by using a hierarchical (multi-level) model to account for the genealogical relatedness of languages on a large and diverse sample of languages (Moran 2012). They find no support for population size as an explanatory predictor of phoneme inventory size, once the phylogenetic relatedness of languages is taken into account. As such, there is currently no support for a positive correlation between population size and phoneme inventory size --- which is crucial to Atkinson’s model.

Moreover, the use of current population counts is also problematic for several reasons. First, when modern humans left Africa, Pleistocene populations would have been small and subdivided; languages would have had, perhaps, 100,000 speakers at most (Sproat 2011; Ringe 2011). Estimates for the population of humans around 10kya before present are roughly one million, so that Atkinson’s time scale of 50-70kya would translate to even smaller population figures. Therefore, the correlation put forth by Hay & Bauer (2007) is mute because population ranges in the Paleolithic are not necessarily reflected in today’s population statistics.^4^ Large community sizes arose only after the peopling of the entire world and the adoption of agriculture. Moreover, synchronic population data is not necessarily reflective of the long-term socio-cultural history of speaker communities (Dahl 2011). These facts also provide points of contention for skeptics of Atkinson’s results.

For example, Atkinson would have to minimally show that Hay & Bauer’s (2007) finding holds over population ranges from a few speakers to a few thousand speakers (Liberman 2011). However, Cysouw et al. (2012) show that contrary to Atkinson’s findings, the correlation between population size and phoneme inventory size only reaches significance when languages with speaker populations above 10^5^ are included.

Paleolithic languages would have been spoken by relatively small populations (Bowern 2011). Sproat (2011) asks if there is any reason to believe that a population of, say, tens of individuals would show less linguistic diversity than their founder population which may have consisted of, at most, hundreds of individuals? (Issues regarding language transmission are discussed below.) Simply put, there is no evidence that current population figures are reflective, or even plausible, of the Paleolithic (Sproat 2011; Liberman 2011). And without the positive correlation between population size and phonemic diversity, Atkinson’s conclusions do not hold.

A second point of scrutiny is the fact that colonization in recent centuries, as well as migrations in prehistory, make current population figures irrelevant for the time scale of Atkinson’s claims (Bowern 2010; Dahl 2011; Maddieson et al. 2011; Sproat 2011). For example, Liberman (2011) notes that the rate of phonetic innovation and phonemic splits in recorded history makes it implausible that phonological inventory sizes still reflect a serial founder effect in the timescale that Atkinson is investigating. Wichmann et al. (2011) state that it is doubtful that phoneme inventory sizes change at a rate in which they can stay in tune with population sizes given what is known about the rate of sound change and population size increases.

#### 3.2.3. Contradictory points of language’s origin

Atkinson fits his origin model 2560 times (each time considering an origin in a different geographical location) in order to identify the best fitting model. Jaeger et al. (2011) note that the distance effect may not hold up to further testing and that there is a high chance to find a significant distance effect, even if there is none (thus, Atkinson’s approach has a high Type I error rate). Although Jaeger et al. (2011) find the best fit for a single origin model lies in West Africa, after taking geographic effects into account, and thus replicating thus Atkinson’s results with Atkinson’s data -- other studies did find contradictory points for the origin of language using similar approaches.

Wang et al. (2012) show that the WALS simplification of phoneme inventory counts distorts Atkinson’s results and the decline in phonemic diversity is not obtained when the data are not binned (cf. 3.1 above). When using count data, they find the most appropriate best-fit origin for modern languages using Atkinson’s method is Asia.

Cysouw et al. (2012) show that phoneme inventory size in WALS is imperfectly correlated with the number of phonemes reported in UPSID (r = 0.60; cf. 3.1 above). This leads to a strong bias in geographic patterning in Western Africa in Atkinson’s results due to the WALS data giving unjustified weights to the number of vowels and tones at the expense of consonants (as discussed above). Cysouw et al. (2012) find that when the UPSID data are corrected for population size and linguistic genealogy in a multi-level model, the largest phoneme inventories are actually found in North America. After replicating Atkinson’s method with the UPSID data, Cysouw et al. (2012) find two equally well supported, but separate points of origin: one in Eastern Africa and one in the Caucasus.

Van Tuyl & Pereltsvaig (2012) show that the point of origin changes negligibly when the correlation is maximized by using only African data. They demonstrate that the non-African data are irrelevant for determining the point of maximum phoneme diversity within Africa. Therefore, they suggest an extrapolated point of origin with maximum correlation may be an artifact of Atkinson’s analysis.

Lastly, we note that the use of modern latitude and longitude coordinates for languages may not be reflective about where their ancestral languages were spoken in the past. Ancestral heartlands are not necessarily taken into account when geo-coordinates are compiled from various sources, such as grammars and sociolinguistic surveys, because the histories of languages and language families is often unknown.

#### 3.2.4. The processes involved in how phoneme inventories change

A phoneme is a unit of analysis, and as already pointed out, different methods of analysis will yield different numbers of phonemes (examples are given in Donohue & Nichols (2011), Maddieson et al. (2011), and Rice (2011)). A phoneme is an abstraction of some number of phonetic variants that are perceived as equivalent by speakers of a language. It is not the case that a subgroup of speakers can be expected to use only a subset of the phonemes in their language (Maddieson et al. 2011).

Therefore, any “diversity” lost to migration would be phonetic, not phonemic, due to the reduction of contact among the number of dialect speakers. This means that the relevant level for diversity reduction is at the phonetic level, not the phonemic one (Bowern 2011; Maddieson et al. 2011; Pericliev 2011). As such, Bybee (2011) suggests that looking for causal mechanisms that relate the number of phonemes in an inventory to population size requires looking at sound change. However, the relationship between sound change processes and the number of phonemes that are created or lost is indirect and whether sound change itself relates to population size is questionable (Bybee 2011). For example, if small populations may exhibit slower rates of sound change due to their dense social networks (Bowern 2011), then the founder population would have fewer sound changes with which to maintain their phonemes (Bybee 2011). But phonological change is independent of human migration and it follows certain pathways driven by production and perception pressures. As such, sound change causes both increases and decreases in the number of phonemes, and the number of phonemes is ultimately a property of the lexicon (Maddieson et al. 2011). The loss of a phoneme would involve the loss of particular lexical items containing that phoneme by the founder population, which seems highly unlikely given what we know about phonological change:

For example, it is unlikely that phoneme inventories bear a signal from the distant past (in Atkinson’s model 50-70kya) due to the fact that phonological change takes place too quickly and too often for phoneme inventory sizes to reflect ancestral states, say, even 10kya (Bybee 2011). For example, Bybee (2011) notes that Latin’s descendant languages, including French, Spanish, and Portuguese, bear little similarity with one another in terms of vowels, phonotactic patterns, rhythmic type, etc. Each has more consonants than Latin, despite radiation out of Rome.

Liberman (2011) also notes that there are many examples of parent languages that give rise to descendent languages with larger phoneme inventories and phonemic diversity. One example is Bantu, in which Proto-Bantu is reconstructed as having 25 phonemes, but many of its descendants have larger inventories due to borrowing click speech sounds from Khoisan (e.g. Xhosa, Sotho, Zulu). As such, modern African languages may have larger inventories than their ancestors, which would overestimate phoneme inventory sizes in the Paleolithic. On the other hand, and as already discussed, Austronesian languages have decreased in their number of phonemes over the last several thousand years as humans migrated throughout Oceania (Trudgill 2004). Hence, given that languages in relatively short time spans can undergo significant changes in their sound systems, Liberman (2011) notes it is implausible that Atkinson’s method has identified a phonological signal that is as old as 50-70kya.

Lastly, if geographic isolation is taken as a proxy for bottleneck effects, this does not necessarily lead to the loss of phonemes in languages. Clearly, human migration within geographic areas may shape phonological inventories because the sounds of a language are transmitted from parents to children, because languages may come into contact (Thomason 2001), and because sound systems are affected by various environmental and climatic variables (Everett, Blasi, and Roberts 2016; Dediu, Janssen, and Moisik 2017), but human expansion out of Africa has left no strong signature on phonological inventories (Creanza et al. 2015).

Taken together, these considerations cast doubt on any causal mechanism behind Atkinson’s conjecture. This stands in marked contrast to the explicitly tested causal mechanism that we develop for the correlation between labiodentals and subsistence.

#### 3.2.5. No theoretical or empirical support for a serial founder model for language

Despite the issues raised so far regarding suboptimal data, the lack of a correlation between population size and phoneme inventory size, different points of origin, and issues in the underlying mechanisms of change, T&U note that Jaeger et al. (2011) largely replicate the results of Atkinson’s (2011) statistical methods after taking language contact and genetic dependencies into account. However, Jaeger et al. (2011) address two serious problems with the findings by Atkinson, which leads them to ultimately conclude that there is no strong support for Atkinson’s serial founder model, noting also that Cysouw et al. (2012) were unable to replicate Atkinson’s results on an alternative dataset.

First, Jaeger et al. (2011) find the Type I error rate is much higher in Atkinson’s analysis than conventionally accepted. Second, their model evaluation suggests that although most assumptions of Atkinson’s model are met, there are potential problems with the assumption of homoscedasticity. In a model with random intercepts for language genealogy (family, subfamily, genus), many grouping factors are represented by single languages. How can one then reliably assess whether the residuals exhibit the same variance for the grouping factors at all levels (Jaeger et al. 2011, Appendix B)? Ultimately, Jaeger et al. (2011) show considerable between-language family variation in regard to effect from distance from Africa. However, Atkinson’s findings are only supported if the sampled languages for small languages families are considered to be representative for the entire language family. This assumption remains untested.

## 4. Poisson Linear Regression (PLR): model comparison

Our own study shows that models where subsistence plays a role through interactions can still provide a good fit to the data in comparison to the model without subsistence at all (Figure S3 in the supplementary information of our paper). This is not surprising to the extent that the effects of subsistence on diet and bite will be strongly modulated by region and family. Finding an effect at the level of interactions is in line with the conditional model we spelled out before: once all other sources of labiodentals are taken into account, there is a detectable effect which could still differ according to the levels of the conditioning variables. In any case, we want to point out that our state-of-the-art model evaluation approach still finds more support for the model with a main fixed effect for subsistence in contrast to the one where it only features in interactions.

## 5. Phylogenetic Analyses

We found the phylogenetic analyses proposed by T&U intriguing, and, while we think they are problematic for several theoretical reasons described in detail below, we nevertheless tested them by re-implementing their method using the software of their choice (`RevBayes`, which is different from what we used in our paper).

### 5.1. Theoretical objections

T&U suggest that the increase in labiodentals in Indo-European could simply be a reflex of the stochastics of Markov processes, in which case the phylogenetic models we used in the paper would not be able to falsify our hypothesis. This would indeed be a challenge if all cognate sets would evolve with the same transition rates and would have stabilized within the lifespan of Indo-European. But as Fig R5 shows, these rates differ greatly. Our hypothesis would be falsified if all cognate sets behaved like *b^h^ or *p, which start with low probabilities and end up with higher probabilities. But there are also sets like *k^j^, which developed labiodentals only in English and Albanian (e.g. in the English word “enough”), or *m, which changed back and forth multiple times (as shown in our paper), without stabilizing. The identification of characters like *k^j^ that keep low probabilities throughout the lifespan of Indo-European is consistent with findings from other studies of language change using Bayesian phylogenetic inference: for example, bans on recursion in syntax never gained much probability in Indo-European (Widmer et al. 2017).

**Figure R5:**
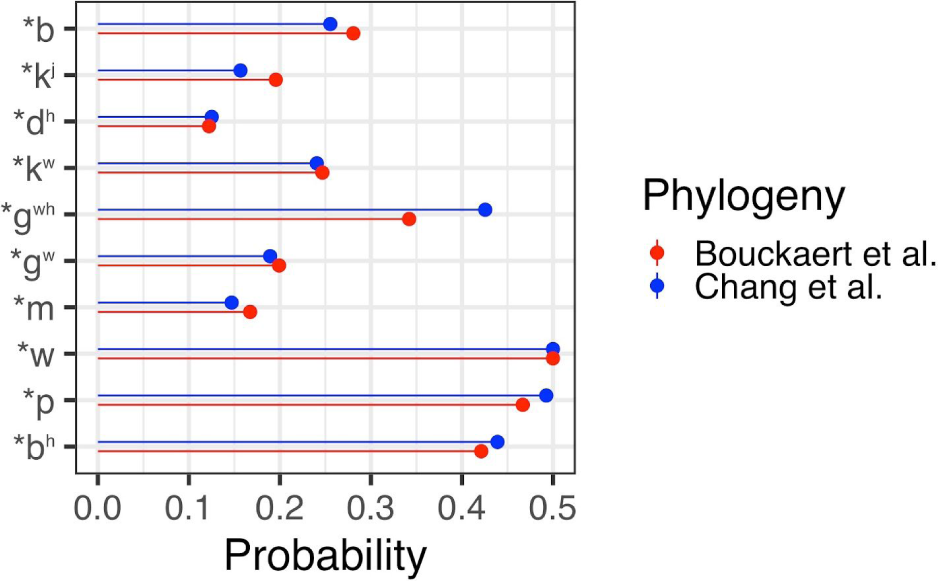
Probabilities of gaining a labiodental after 8,000 years, according to the models fitted in the original paper. The characters (the cognate sets, represented by the rows) behave very differently.

T&U fail to recover the historical heterogeneity among branches that our model suggests (Figure 7B in our original paper), but this is clearly at odds with attested developments, such as in Indo-Iranian vs the European clades. This raises the possibility that T&U’s result is in fact a *null result* and therefore not evidence against branch heterogeneity (see also below). As we note in our paper, several confounds contribute to the heterogeneity that we observe (e.g. the attested developments in Indo-Aryan, Greek or Romance), including the historical dynamics of contact within and outside of Indo-European, different language ideologies (conservative vs innovative attitudes), and the social dynamics (prestige vs non-prestige status of adopters of sound change). All of these factors are likely to mask branch-specific developments, and we would be surprised if heterogeneity could be modeled without controlling for them. For this reason, we refrained from such models in our original study and limited ourselves to note the overall heterogeneity that emerged from Stochastic Character Mapping models without any prior constraints on regional or cladistic differentiation. In this regard, our results are consistent with attested history.

Another important concern is that T&U collapse the cognate sets. This is at odds with how sound change proceeds: it is not a sampling process among just any kind of sound. Instead, sound change is a sampling process among the phonetic variants (allophones) of each *cognate* sound across speakers and this is represented in each language as a phoneme, a functionally distinctive unit (Blevins 2004; Ringe and Eska 2013) in a language. For example in English, the phoneme “p” has two allophone variants, the plain form [p] which occurs only after sibilants, as in [spin] and the form [p^h^], as in “pin”, which occurs everywhere else. This tie to phonemes is the reason why sound change is regular and constrained by the system of contrasts in each language and *must* be modelled as such.

### 5.2. Technical issues

We started from the `R` and `RevBayes` scripts that T&U have made available in their GitHub repository,^5^ and we adapted, streamlined, improved and heavily commented their code in order to make it easier to test alternative models and variations thereof without overloading the user with several individual scripts (now the scripts can be customized from, for example, `BASH`, using parameters). We have made available these customizable `RevBayes` and `BASH` scripts that use it to fit the various models and variants described below, as well as a massive `Rmarkdown` script and resulting `HTML` document that contains all the necessary plots, analyses and results referred to here. These are available in our GitHub repository. ^**6**^ As such, we only offer here a very brief summary, referring readers to our full analysis report in the GitHub repository.

First, we observe that the T&U `RevBayes` script is actually based on a `RevBayes` tutorial (`m_DPP_bears.Rev`, available at: (https://raw.githubusercontent.com/revbayes/revbayes_tutorial/master/RB_DPPRelaxedClock_Tutorial/scripts/m_DPP_bears.Rev) which implements the *Dirichlet Process Prior* (DPP) model that they use. However, we also observe that, for no clear reason that we can identify, they removed part of this tutorial script, which implements an overall *base rate* (a real number with a log-normal prior that is multiplied with the DPP branch rates to produce the actual branch rates of the tree) -- we refer to this (“reverted” to the tutorial) model as **TUBR** (<u>T&U</u> with <u>B</u>ase <u>R</u>ate).

Our second observation is that the DPP process makes `RevBayes`’ mechanism for estimating the marginal likelihood (and, more precisely, the `powerPosterior.run()` function) fail,^7^ which means that *Bayes Factors* (BFs) cannot be computed for the original **TU** model, nor for its “reverted” variant **TUBR**, making formal model comparisons involving them not possible (this may be the reason for which they only used visual comparison in `Tracer` for judging model fit in their comment).

Our third observation concerns the mechanism used by T&U to sample the individual trees from the full tree sample that we used in our paper: this is based on an integer variable that can take values between 1 and 1000 (they limit themselves to a 1000-trees subsample of our full sample, probably for computational reasons) and which represents the tree in the sample currently used. However, while this mechanism works well for the **TU** model (in the sense that pretty much the whole sample is visited during the MCMC process), for the others it does not, and instead it becomes “stuck”, with just a few (sometimes only 1) tree(s) being visited, despite the much larger number of generations and burn-in we used. We think what happens is that, while the **TU** model fits uniformly bad (see below) all trees in the sample, the other models actually fit individual trees with different parameters, making the jump to a different tree (once the other parameters are optimised for the current one) very improbable. As a consequence, besides using T&U’s mechanism (a), we also implemented (b) a fit using only the *MCC consensus tree* and (c) *separate fits* for as many individual trees from the sample as our computational resources allowed for this response. Our final interpretations are based on all three of these considerations (see the full `HTML` report in our GitHub repository).

Because of these issues -- coupled with the theoretical problems discussed in Section 5.1 above -- we also implemented a set of *alternative models and variants*, as follows (see the *Data, Tree sampling, Implementation, Models* and *Methodology* sections of the `HTML` analysis report on GitHub for full details):^8^

- <u>Models</u>: besides **TU** and its “reverted” base-rate version, **TUBR**, we implemented also the *uncorrelated branch rate* (**ULR**), the *molecular clock* (**MC**) and the *relaxed molecular clock* (**RMC**) models,
- <u>Variants</u>: for the **root**, we implemented either the original TU model where these come from the stationary distribution (denoted **1**), and a variant where they come from a *Dirichlet distribution* (denoted **DP**); for the **sites** (i.e., the characters), we implemented either the original TU single-rate across characters model (denoted **1**), or a *mixture model* with two site classes (slow and fast; denoted **MM**).

Thus, we ended up with 12 model variants in total (TU-1-1 [the original TU], TU-DP-1, TU-1-MM, TUBR-1-1, ULR-1-1, ULR-DP-MM, MC-1-1, MC-DP-MM, RMC-1-1, RMC-DP-1, RMC-1-MM, RMC-DP-MM), which cover a large space of possible models, and allow us to estimate **TU**’s robustness to various changes. As mentioned above, except for **TU** and **TUBR**, we could use Bayes Factors (BFs) for model comparison, while for comparisons involving these two we were forced to use only the (log)likelihoods.

With these, our main results (in brief) are:

1. the **TU** model *does not fit the data*, being pretty much the worst among the models and variants we tested;
2. the **TU**’s estimate of the number of *branch rate categories* -- considered by T&U as the relevant measure in their response -- is *deviant* compared to the other models: it is the only model that produces 1 category (apparently, this is a constant with no variation due to the MCMC process, which is relatively surprising), while the other models do produce distributions of estimates (as expected from a Bayesian model) centered around values larger than 1 (for **TUBR**, this is centered around 2);
3. the *values estimated at the root* are also *deviant* for the **TU** model (50%:50% for presence:absence), but are much more in line with our previous estimates (around 0% presence for most models, including **TUBR**) and consistent across models.

Thus, the **TU** model’s results are *not* robust, it *fits very badly the data* compared to the other models (even compared with such an unrealistic model as a single-rate molecular clock), and even adding back the base rate (a simple numeric multiplier) makes it produce estimates in line with the other models and with our own results in the paper. This applies to the *root value estimates* (extremely low probability of presence in Proto-Indo-European), as well as to the *number of rate categories* (definitely higher than 1, its actual distribution depending on the model).

## Footnotes

1 Atkinson’s study was also based on faulty WALS data (a misalignment in the input spreadsheet for consonant inventories, which has a marginal impact on the reported results (Maddieson et al. 2011).

2 Ringe (2011) notes the unusual development of phoneme inventories in Oceania may be a language-internal process due to phonotactic constraints, leading to a loss of consonants (Trudgill 2004), and not related to language contact or isolation.

3 The best current estimate for the number of languages spoken as a first language is 7592 (https://glottolog.org/glottolog/glottologinformation).

4 We must also note that most present-day population sizes crucially depend on subsistence methods and cultural and technological practices that were clearly unavailable to the humans migrating out of Africa, and that massively have altered the carrying capacity of various eco-climatic niches.

5 https://github.com/sergeitarasov/Tarasov_Uyeda_SupplementaryMaterials_2019

6 https://github.com/IVS-UZH/response2TU

7 With the very unhelpful error message “Cannot apply Gibbs sampler when the probability is heated”.

8 https://github.com/uzling/response2TU

